# Preliminary cross-sectional investigations into the human glymphatic system using multiple novel non-contrast MRI methods

**DOI:** 10.1101/2023.08.28.555150

**Authors:** Swati Rane Levendovszky, Jaqueline Flores, Elaine R. Peskind, Lena Václavů, Matthias J.P. van Osch, Jeffrey Iliff

## Abstract

We discuss two potential non-invasive MRI methods to cross-sectionally study two distinct facets of the glymphatic system and its association with sleep and aging.

We apply diffusion-based intravoxel incoherent motion (IVIM) imaging to evaluate pseudodiffusion coefficient, *D\**, or cerebrospinal fluid (CSF) movement across large spaces like the subarachnoid space (SAS). We also performed perfusion-based multi-echo, Hadamard encoded multi-delay arterial spin labeling (ASL) to evaluate whole brain cortical cerebral blood flow (CBF) and transendothelial exchange (T_ex_) of water from the vasculature into the perivascular space and parenchyma. Both methods were used in young adults (N=9, 6F, 23±3 years old) in the setting of sleep and sleep deprivation. To study aging, 10 older adults, (6F, 67±3 years old) were imaged after a night of normal sleep only and compared with the young adults. *D\** in SAS was significantly (p<0.05) lesser after sleep deprivation (0.014±0.001 mm^2^/s) than after normal sleep (0.016±0.001 mm^2^/s), but was unchanged with aging. Cortical CBF and T_ex_ on the other hand, were unchanged after sleep deprivation but were significantly lower in older adults (37±3 ml/100g/min, 476±66 ms) than young adults (42±2 ml/100g/min, 624±66 ms). IVIM was thus, sensitive to sleep physiology and multi-echo, multi-delay ASL was sensitive to aging.

## INTRODUCTION

The glymphatic system is a brain-wide network of perivascular pathways along which subarachnoid cerebrospinal fluid (CSF) and brain interstitial fluid exchange, supporting solute distribution and clearance^1–3^. Implicated in the clearance of amyloid-β^2–4^, tau^5,6^, and α- synuclein^7,8^, impairment of glymphatic function is proposed to contribute to the development of neurodegenerative dementing disorders such as Alzheimer’s disease. Initial studies in mice characterized perivascular glymphatic function in mice using dynamic 2-photon microscopy following intrathecal fluorescent CSF tracer injection^2^, an approach with high spatial and temporal resolution, but with a limited dynamic field of view. Brain-wide glymphatic exchange, organized around the macroscopic cerebral vasculature, was subsequently described in rodents using dynamic contrast-enhanced magnetic resonance imaging (MRI) following intrathecal gadolinium-based contrast agent (GBCA) administration^1^.

Studies by Ringstad, Eide and colleagues have used a similar intrathecal approach to confirm key features of the glymphatic transport model that was initially described in rodents, in the human brain. In ‘reference’ participants being evaluated for neurosurgical intervention, these authors report that serial MRI scanning following intrathecal GBCA injection results in initial cisternal and subarachnoid space (SAS) enhancement, followed by a centripetal enhancement of the gray and white matter tissue^9^. Using serial T1 mapping over three days, Watts et al. showed that signal enhancement peaked in the CSF and gray matter (GM) at approximately 10 hours and in white matter (WM) at approximately 40 hours^10^. Sleep-active solute clearance from the human brain was also confirmed in a recent study, comparing the effects of overnight sleep deprivation on the clearance of intrathecally-injected GBCA^11^. Since intrathecal injections are an off-label use for GBCA and difficult to implement in a research setting, several recent studies have sought to extract similar information from MRI following intravenous GBCA injection. In one study, heavily T2-weighted fluid-attenuated inversion recovery (FLAIR) showed GBCA transport from the vasculature into the CSF via the choroid plexus, along perivascular spaces and perineural sheaths of cranial nerves. In a recent study, we further characterized CSF, GM, and WM enhancement that occurs between 0-6 hrs. following IV GBCA administration^12^. While these contrast-based methods for assessing solute transport through the CSF and brain interstitium remain a gold standard, these approaches remain technically challenging and suffer from poor temporal resolution. Functional MRI studies revealed that coordinated CSF pulsations increase during sleep, perhaps reflecting increased activity of the glymphatic system CSF ^13^. These coordinated pulsations appear to arise from slow wave-associated low-frequency vasomotor fluctuations that are hypothesized to be one of the drivers of perivascular glymphatic convective transport of solutes^14,15^.

Here we present two modified diffusion and perfusion MRI methods as potential markers of glymphatic physiology: (1) Intravoxel incoherent motion (IVIM) MRI, a diffusion-based approach differentiating water mobility into different components like perfusion, water diffusion in tissue, interstitial fluid (ISF) and CSF, and (2) multi-echo Hadamard encoded arterial spin labeling (ASL) to study water exchange from the blood compartment into the brain interstitium. IVIM conceptually detects a second diffusion regime in the tissue at lower b-values that corresponds to a slower movement over long distances (also known as a faster pseudodiffusion coefficient), *D\**, which was originally proposed to measure perfusion^16^. This is in contrast to the typical slow diffusion coefficient, *D*, measured with diffusion imaging which corresponds to faster movement over very small distances. We believe that the IVIM effect can be exploited within the SAS to characterize CSF mobility, whereas IVIM measurements in the tissue are mainly influenced by perfusion, making it more difficult to e.g. characterize ISF-mobility. We applied an advanced ASL sequence to calculate blood flow and ensure that the *D\** signals are not entirely driven by perfusion. This approach allows us to quantify the trans-vascular exchange of water, which is relevant in the glymphatic transport of solutes. While this measure is influenced by blood-brain barrier permeability, it may also represent changes in glial water transport through aquaporin-4 (AQP4) channels. A study in AQP4 knockout mice showed that trans-vascular exchange increased in the knockout mice by almost 30% when compared to wild-type mice and decreased significantly in aged mice^17^. Only a few studies have explored this advanced ASL approach in humans and in the context of sleep modulation. In the present study we used a within-participant paired sleep/sleep deprivation design to test the ability of IVIM and multi-echo ASL to detect sleep-related changes in CSF-ISF circulation as well as an across participant design to test the sensitivity of IVIM and multi-echo ASL to detect aging related changes in CSF-ISF circulation.

## METHODS

### Participant characteristics and study procedures

This study was approved by the Institutional Review Board of the University of Washington (UW) School of Medicine and all participants provided written informed consent before any study procedures in accordance with the Declaration of Helsinki. Ten normal healthy older adults (6F/4M, age = 67±3 years old) and fifteen normal healthy young adults (6F/7M, age = 23±3 years old) were enrolled. Demographic information is detailed in Table 1. Cognitively normal older adults were identified and referred by the UW Alzheimer’s Disease Research Center (ADRC). They underwent a brief cognitive performance evaluation and a single MRI scan after a night of normal sleep for this study. Cognitive testing included the General Practitioner’s Cognitive Assessment (GPCOG)^18^ and the Memory Impairment Screen (MIS)^19^. Participants were asked to perform their regular activities before coming in for their study visit.

**Table 1:** Parameter values (Average ± Std. Deviation) for IVIM.

| Parameter/Region | Old | Young |  |
| --- | --- | --- | --- |
|  |  | Normal sleep | Sleep deprivation |
| $D^*$ mm<sup>2</sup>/s | | | |
| SAS | 0.016±0.001 | **0.016±0.001** | **0.014±0.001** |
| CP | 0.019±0.001 | 0.019±0.001 | 0.017±0.001 |
| GM | 0.013±0.001 | 0.014±0.001 | 0.014±0.001 |
| WM | 0.013±0.001 | 0.014±0.001 | 0.013±0.001 |
| $D$ mm<sup>2</sup>/s | | | |
| SAS | 0.0014±0.0002 | 0.0013±0.0002 | 0.0012±0.0003 |
| CP | 0.0024±0.0003 | 0.0019±0.0006 | 0.0019±0.0006 |
| GM | 0.0008±0.0001 | 0.0008±0.0004 | 0.0008±0.0005 |
| WM | 0.0007±0.0005 | 0.0007±0.0003 | 0.0007±0.0003 |
| $f$ | | | |
| SAS | 0.17±0.02 | 0.17±0.02 | 0.17±0.02 |
| CP | 0.19±0.04 | 0.21±0.02 | 0.21±0.03 |
| GM | 0.10±0.01 | 0.10±0.01 | 0.10±0.01 |
| WM | 0.09±0.01 | 0.08±0.01 | 0.09±0.01 |

IVIM = intravoxel incoherent motion, SAS = subarachnoid space, CP = choroid plexus, GM = gray matter, WM = white matter

Young, healthy adults were recruited using flyers in and around the institution. They underwent two scans, one after a night of normal sleep and one after 24 hours of sleep deprivation. They were also instructed to follow their regular and identical routines for both visits except for the sleep deprivation. They completed the GPCOG and MIS at each MRI visit. For the sleep deprivation visit, participants were asked to stay awake for 24 hours before their MRI scan. Starting the morning before, they were instructed to complete their normal routine and not nap. From 8 pm onwards, at every hour, they called a designated phone line to let the study team know that they were awake. If they missed a check-in, their scan was canceled. The order of sleep and sleep deprivation visits were randomized. All scans were conducted between 0800- 1000 hours for all participants. For the young adults, both scans were scheduled at the same time in the morning.

### Imaging

All imaging was performed on a 3T Philips Ingenia Elition scanner with a 32-channel head coil. Identical MRI scans were performed on all participants. An eye tracker (Eyelink 1000) was used to check that the participant was awake. In between scan sequences, the technologists or study team spoke to the participants to ensure they did not sleep. The MRI protocol consisted of (1) 3D T1 acquisition with resolution = 1×1×1 mm^3^, matrix size 256×256×176, repetition time (TR)/ echo time (TE) = 9.2/3.5 ms, (2) Intravoxel Incoherent Motion (IVIM) protocol using 12 b-values (10, 20, 40, 80, 100, 150, 200, 300, 500, 700, 900, 1000 s/mm^2^) and 6 directions and one b=0 s/mm^2^, resolution = 1×1×5 mm^3^, slices = 30, TR/TE = 3000/62 ms^16^. To reduce motion-related distortion, a two-echo SPLICE acquisition was used^20^. SPLICE or Split acquisition of fast spin echo approach, modifies the readout to collect two echoes, which are combined to reduce distortion and increase signal-to-noise ratio or SNR. (3) T2-prepared Hadamard encoded arterial spin labeling (ASL) with background suppression and resolution = 3.75×3.75×5 mm^3^, slices = 18, label duration = 3400 ms, effective post labeling delays = 650, 1210 and 2083 ms, and echo time, TE = 0, 40, 80, 120 ms^21,22^. A second reference, M0, scan at all four TE values, was acquired with identical parameters but no labeling or background suppression. All MRI data will be available upon request.

### Analyses

The IVIM MRI was processed to calculate perfusion or fluid fraction, *f*, and pseudodiffusion coefficient, *D\**, as a measure of CSF mobility in the SAS. For this purpose, IVIM data were motion corrected and aligned with the B0 image. Due to SPLICE acquisition, no distortion correction was necessary. Next, the diffusion data was skull-stripped in FSL^23^ and fitted to a two-compartment model

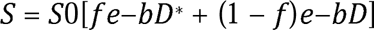

where S0 = signal intensity at b=0 image. *D*, *D\**, and *f* are calculated using Bayesian estimation in MATLAB. The *D*, *D\**, and *f* were then registered to their T1 image in a 2mm MNI space atlas. The T1 image was segmented into GM, WM, and CSF. GM and WM masks were obtained using a probability threshold of 80%. A dilated brain mask was used to eliminate the brain stem regions, and the resultant CSF-mask was thresholded to include only >95% CSF. This stringent threshold was used to ensure there was no partial volume with cortical GM. Due to the partial coverage of IVIM, only CSF space around the frontal cortex was identified as the SAS. All ROI generation was automated (**Supplementary Figure 1**). GM ROI is shown in magenta and overlaid on the participants’ T1 image in MNI space. White matter ROI is shown in white. CP is shown in yellow, and SAS in green.

Cortical GM and subcortical WM were also used as ROIs for comparison while being aware that the IVIM effect could be due to CSF&ISF mobility, perfusion, or a combination of both. Additionally, an ROI was manually drawn in the choroid plexus (CP), representing CSF production and an area showing both perfusion and CSF-related IVIM effects.

The Hadamard-encoded ASL data were rearranged to calculate perfusion-weighted difference maps at the post-labeling delays of 650, 1210, and 2083 ms. The difference maps were registered to the M0. First, the 4-echo M0 data were fit to an exponential decay to estimate extravascular T2 (assuming that most signals were extravascular). Based on the work of Ohene et al., we fit a two-compartment (intravascular and extravascular) model to the difference image at a post-labeling delay of 1210 to obtain T2 of the intravascular compartment^17^. Next, the multi-TI data was used to estimate cerebral blood flow (CBF) and arterial arrival time (δ_a_) using FSL’s BASIL Toolbox, followed by an estimate of the tissue transit time (δ). The difference in the two transit times, T_ex_ was used to measure the trans-vascular water permeability. Since ASL measures perfusion and is typically unsuitable for white matter tissue, measurement was made only in cortical GM. The same ROIs as for the IVIM analysis were used. All data is available upon request.

### Statistical Considerations

A two-way, paired t-test was used to compare the MRI measurements after a night of normal sleep and sleep deprivation in young adults. A t-test assuming homoscedastic distributions (Kolmogorov-Smirnov test for normality, p=0.01) was used to compare MRI measurements between young and old adults. Associations with GPCOG and MIS were tested using a linear regression model of all MRI measures.

## RESULTS

All 10 older adults completed cognitive assessment and MRI sessions. Of the thirteen young adults, nine (6F/3M) completed both visits for sleep and sleep deprivation. Of the 4 participants, who did not complete the study, three were unable to comply with the sleep deprivation protocol, and one participant was removed from the study due to incidental findings affecting study interpretations. As a result, their sleep deprivation visit was canceled.

The GPCOG and MIS measures (averages ± std. deviation) in young adults after a night of normal sleep were 9±1 and 8±0, respectively. Their measures after a night of sleep deprivation were 8±1 and 8±0, respectively. In older adults, the corresponding scores were 9±1 and 8±0. Cognitive test scores were not significantly different between groups or conditions.

The results from IVIM studies are shown in **Figure 1**, with representative images provided for IVIM parameters, *D\**, *D*, and *f* (**Figure 1A**) from one participant. Overall data for *D\**, *D*, and *f* following overnight sleep and overnight sleep deprivation are shown in **Table 1** and graphically in **Figure 1B,D and E**. Using a paired t-test, the *D\** values in the SAS were significantly lower in young adults after sleep deprivation compared to the values after a night of normal sleep (**Figure 1C**, p=0.009). No differences were observed in any other IVIM parameters, *D* (**Figure 1D**) or *f* (**Figure 1E**), or in any other region. No IVIM parameters differed between old and young adults.

**Figure 1:**
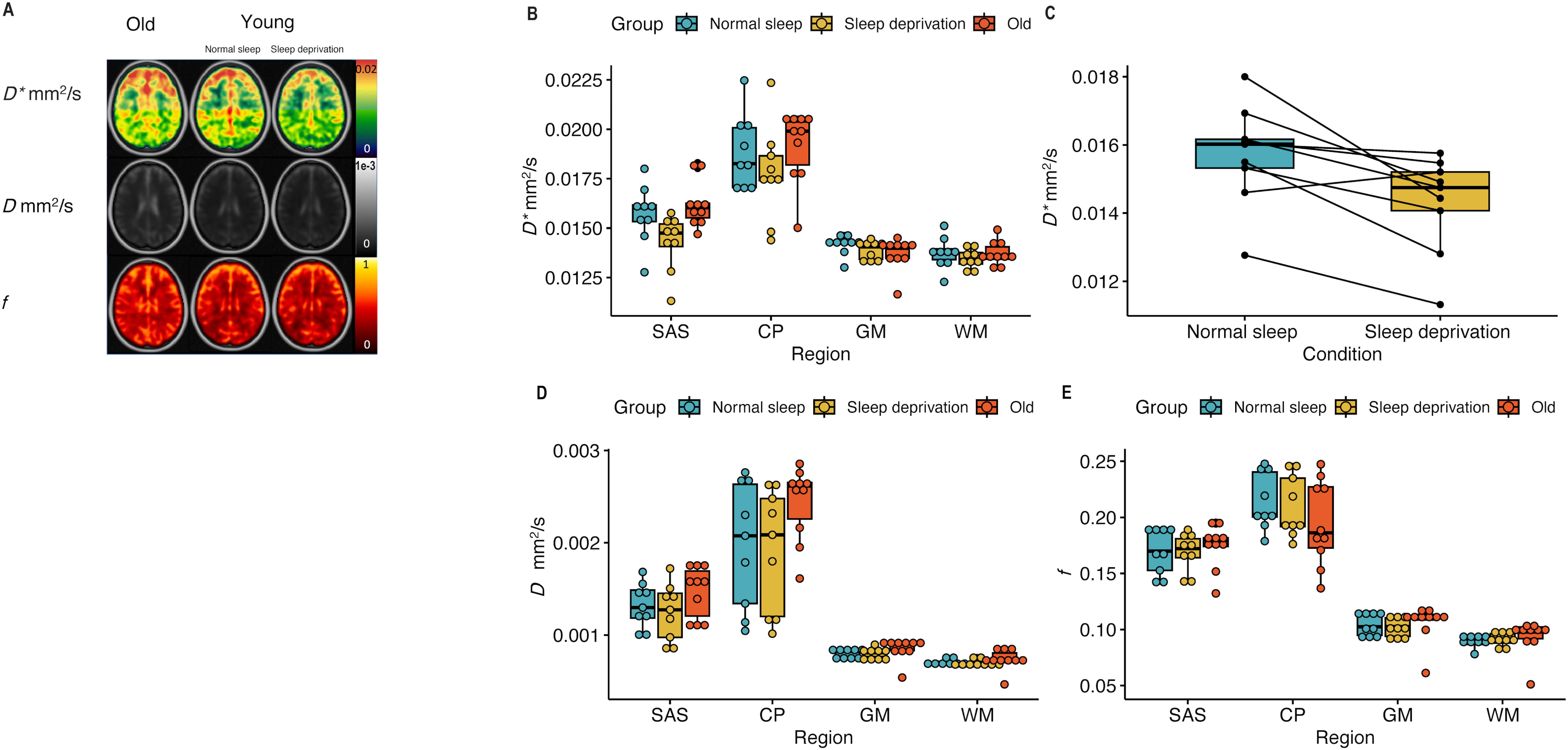
**A.** IVIM parameters, *D*, D,* and *f* in older adults (first column), in young adults with normal sleep (middle column), and the same young adults with sleep deprivation (third column) in the SAS. **B.** *D\** values in all three groups: young adults after a night of normal sleep (blue), young adults after 24 hours of sleep deprivation (yellow), and older adults after normal sleep (red). **C.** Only *D\** was significantly different (p=0.009) after 24 hours of sleep deprivation compared to a night of normal sleep. **D, E.** *D,* and *f* values in all three groups: young adults after a night of normal sleep (blue), young adults after 24 hours of sleep deprivation (yellow), and older adults after normal sleep (red). No difference was observed with aging. **Supplementary figure 1** shows the ROIs generated for this study.

The results from the multi-echo ASL studies are shown in **Figure 2A**, with representative images provided for ASL parameters, T_ex_ and CBF. Overall ASL measures for each group are provided in **Table 2** and visualized in **Figure 2B-C**. **Figures 2B, C** show a comparison in young adults after normal sleep and sleep deprivation and older adults. No significant difference was observed in these measurements. However, there is significantly reduced CBF and T_ex_ in older adults compared to young adults after normal sleep (p=0.005 and p=0.0001, respectively).

**Figure 2:**
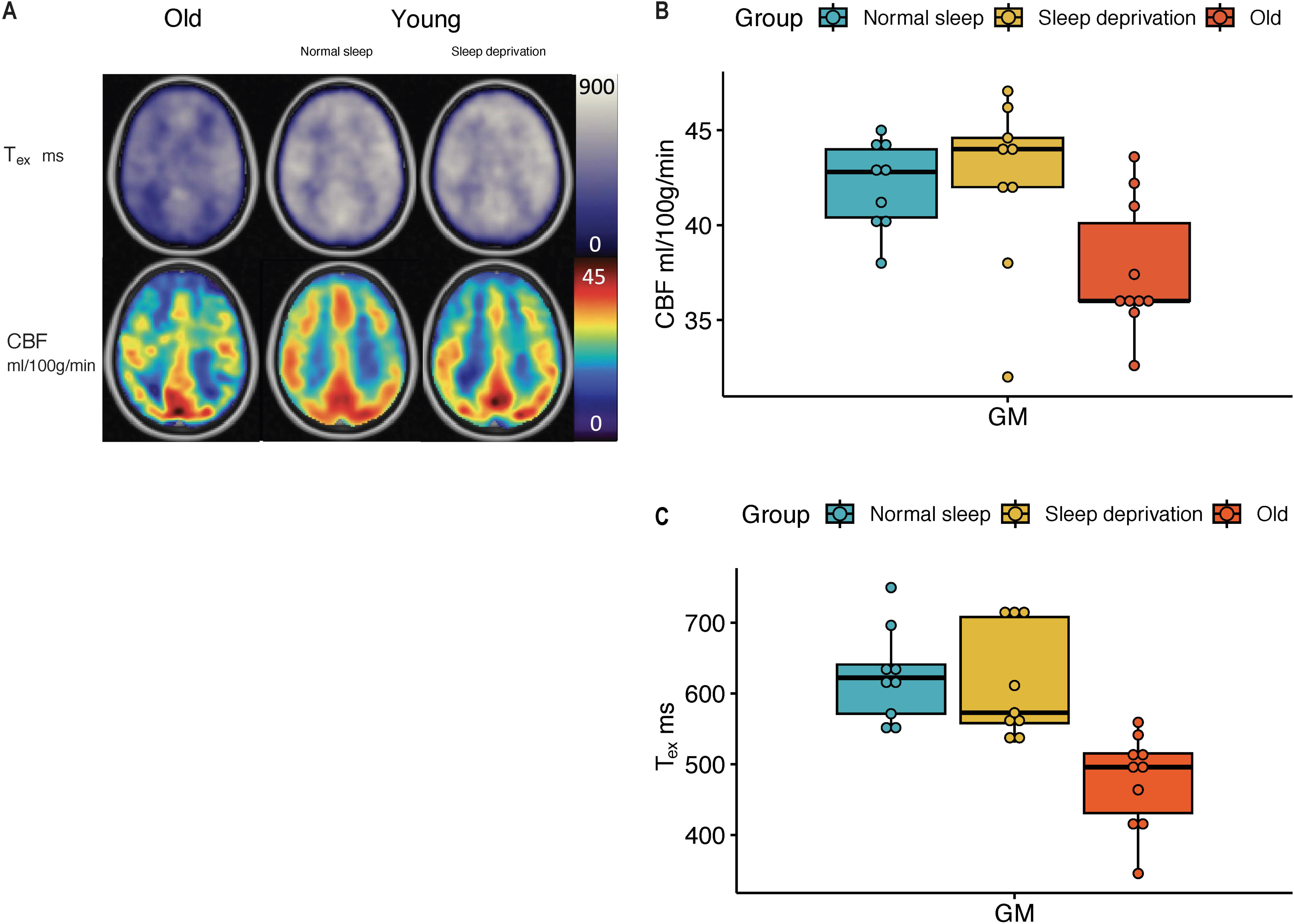
**A.** ASL parameters, Whole-brain cortical CBF and T_ex_ in older adults (first column), in young adults with normal sleep (middle column), and the same young adults with sleep deprivation (third column). CBF was not significantly different in young adults after sleep deprivation (yellow), when compared to those after a night of normal sleep (blue). Detailed paired comparison is shown in **B.** CBF was significantly different in older adults (red) compared to young adults (p=0.005). **C.** T_ex_ was also not significantly different in young adults after sleep deprivation (yellow), when compared to those after a night of normal sleep (blue). Like CBF, T_ex_ was significantly shorter in older adults (red) compared to young adults (p=0.0001).

**Table 2:**
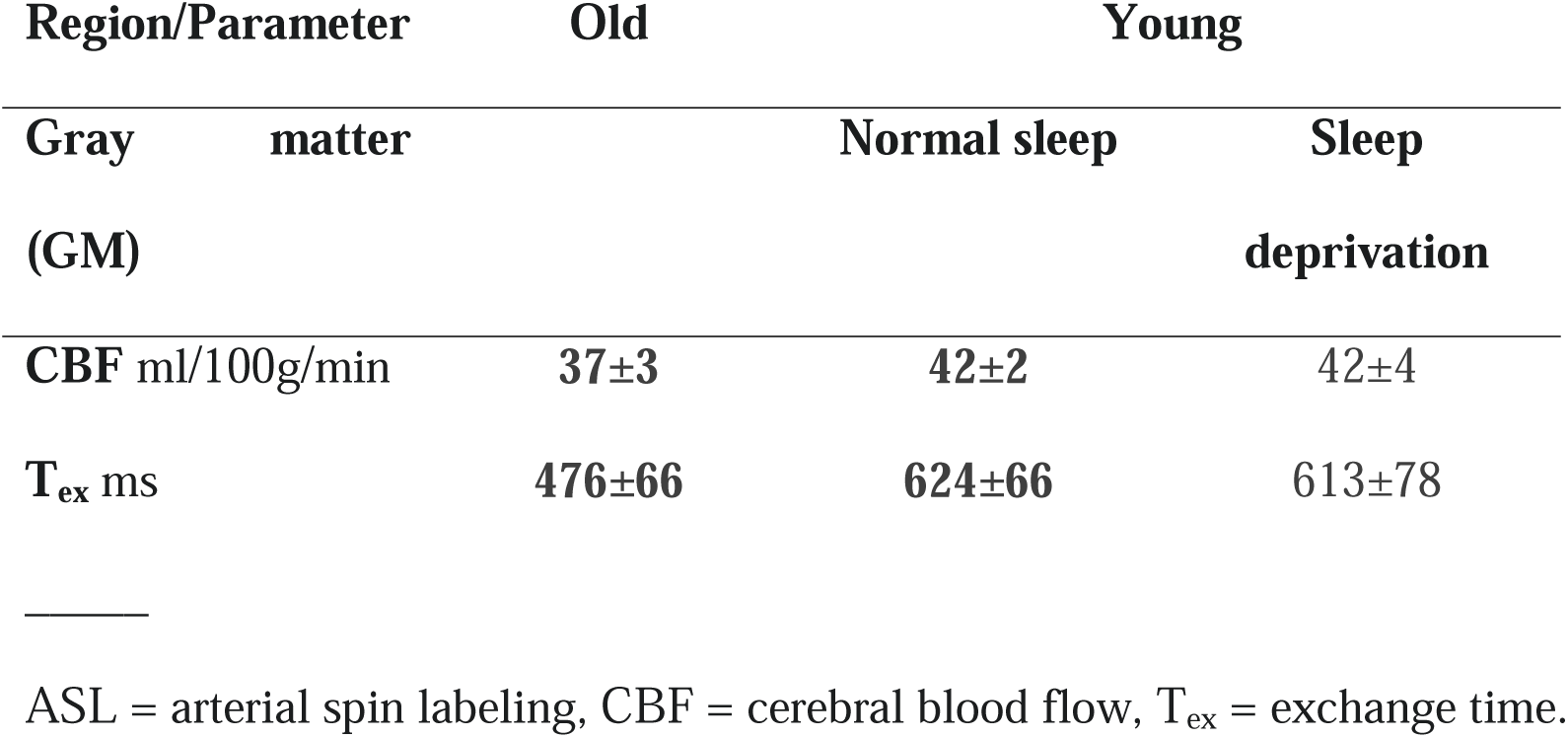
Parameter values (Average ± Std. Deviation) for ASL.

No significant associations were detected with GPCOG or MIS. However, a lower T_ex_ after sleep deprivation was associated with better GPCOG performance in the young adults (r = -0.70, p=0.04). No other association was significant.

## DISCUSSION

This work explored two novel MRI approaches and their ability to detect aging and short-term sleep-related changes in factors affecting glymphatic transport. Using IVIM, we showed D*, measuring CSF mobility in the SAS, was significantly reduced after sleep deprivation compared to following a night of normal sleep. This parameter, however, was unchanged in healthy older adults when compared to healthy young adults. Using multi-echo ASL, we observed that perfusion and trans-vascular water exchange time were reduced in older adults compared to young adults, while this parameter was not sensitive to overnight sleep deprivation.

A key component of glymphatic transport is the movement of CSF within the SAS space and along perivascular spaces into the brain interstitium. Current approaches using intrathecal gadolinium are difficult to implement for research purposes and unsuitable for repeated measurements to assess changes in glymphatic transport across different physiological or pathological states. Here we used IVIM as an MRI marker of CSF mobility through the SAS as an indicator of glymphatic transport. Le Bihan proposed IVIM as an approach to discriminate microvascular blood flow from water diffusion in the brain, whereas in the SAS, *D\** will reflect slow CSF mobility over large distances as it flows through and around the trabeculae. This SAS IVIM pseudodiffusion effect or *D\**, although not diffusion in the strict sense of the term, represents movement of fluid (i.e., CSF) through the trabeculae within the SAS^24,25^, giving the appearance of random/Gaussian motion of water molecules of the CSF within a voxel as they navigate the trabecular columns in the SAS. IVIM utilizes low b-values with only 3 orthogonal directions of diffusion gradients to sensitize itself to this slow CSF velocity. Low b-values have also been used to assess CSF flow by other studies in different ways prior to our study. For instance, Wells et al. used b = 100 s/mm^2^ to determine the diffusion and direction of CSF flow along the perivascular space of large arteries in rodents^26^. Le Bihan et al. attempted b = 250 s/mm^2^ to determine the s-index to evaluate brain fluid transport, assuming that all perfusion-related IVIM effects are minimized at this value^16,27,28^. However, CSF flow velocity is unknown and likely to vary in the SAS, in the MRI visible perivascular spaces, and around large blood vessels and could span a range of velocities. We did not assume fluid flow velocity cut-offs and use the entire range of b-values from 0 to 1000 s/mm^2^. This imaging configuration makes IVIM measures sensitive to perfusion-related IVIM effects, and the SAS includes penetrating vasculature. We therefore tested in 4 individuals an identical IVIM protocol (with 6 slices) as above, in which signals from all tissue types including GM, WM, CSF, and blood will contribute to the total signal, and compared it with a similar IVIM protocol albeit with prolonged TE (340 ms) to minimize contributions of GM, WM, and blood, i.e. only CSF signals remain. There were very little differences in *D, D*,* and *f* between the two protocols (-3±10%, 2±10%, and 3±4%, respectively). As a validation, we found that in the WM, *D* reduced by 65±10%, *D\** increased by 33±17%, and *f* reduced by 22±9%. As expected, the decreases in *D* and *f* are substantial in the WM. Increases in *D\** are unlikely, and we attribute these increases to fitting inaccuracies in the presence of very low signal in the WM. Interestingly, the IVIM parameters are not substantially changed in the SAS, indicating that *D, D*,* and *f* in the SAS are subject to only minimal contamination by perfusion effects.

IVIM provides three parameters: diffusion, *D,* pseudodiffusion *D\**, and fluid volume *f*. For efficient glymphatic transport, both fluid volume and fluid movement are critical. Due to the high variability and noise in the IVIM signal, the product *fD\** is often used as a measure of fluid movement instead of *D\** alone. We chose to keep the two separate and have identified that with sleep deprivation, *D\** is significantly reduced, but not *f*. Although not observed, it would have been likely that fluid volume changes would occur in the SAS across these conditions. Studies to date observed an increase in interstitial fluid volume with sleep, which in this case would be defined by *f* in the GM and WM^3^. However, the subjects were not scanned during sleep, but after sleep-deprivation which might explain the absence of a clear change in *f*. One drawback of our approach is that in GM and WM we cannot separate the contribution of the perfusion and the CSF compartments.

Many studies have shown that perfusion is reduced with age^29,30^ and in older adults with poor sleep (duration and quality)^31^. However, the acute effects of sleep deprivation are still unknown. A recent study with electroencephalogram (EEG) and ASL showed that non-REM sleep and greater slow wave activity was associated with reduced prefrontal CBF^32^. In our study, we observed aging-related changes in CBF as expected, but we did not observe any change in CBF upon sleep deprivation. These results may be in conflict with prior studies. Zhou et al. recently showed that with 36 hours of sleep deprivation, hippocampal and prefrontal CBF reduced compared to a baseline scan after normal sleep. Another study by Poudel et al. showed that CBF was reduced after 24 hours of sleep restriction but with restricted sleep of 4 hours prior to the restriction. One possible explanation may be that a single night of sleep deprivation (∼24 hours) may not be sufficient to evoke such changes in resting CBF. A second possibility is that regional effects on CBF occur in response to sleep deprivation, but that these effects are obscured by the present approaches that assess global changes in CBF.

The advantage of our ASL approach was that we could measure trans-vascular exchange time or the permeability of the cerebral vasculature to water. Prior studies have shown this measure to be sensitive to aging and *Aqp4* gene deletion in humans and mice^17,33^. No previous studies have been conducted in humans or mice to examine the relationship between trans-vascular water permeability and sleep. Our study found that the permeability, measured in terms of exchange time (T_ex_), was significantly lower with increasing age. The exchange time was also numerically lower after sleep deprivation, but these effects were not significant. This may be because of the relatively small sample size in the present study, or it may be that regional differences in T_ex_ following sleep or sleep deprivation may be obscured using the present global measures. To keep acquisition times low, we used only three delay times in our current implementation which could limit the ability to detect subtle differences in T_ex_. Using more post labeling delays would improve the accuracy of the arterial arrival times and tissue transit times needed for calculation of T_ex_. A different approach to measuring trans-vascular permeability is the use of diffusion-weighted ASL. Gold et al. showed that with this approach, the water exchange rate was faster with aging in humans^34,35^. Our observation of a lower T_ex_ aligns with this previous observation. Diffusion-weighted ASL has not been used to study acute sleep deprivation, but a faster exchange rate was observed in individuals with obstructive sleep apnea^36^. Taken together, we believe that perfusion is likely sensitive to sleep deprivation but may require either a more intense restriction protocol, or greater regional resolution. Both sequences, the multi-echo ASL and diffusion-weighted ASL, have low SNR (SNR being slightly higher in the diffusion-weighted ASL), are non-standard sequences, and availability is vendor and scanner dependent.

Finally, many MRI methods have been proposed to study the perivascular glymphatic transport. Diffusion tensor image analysis along the perivascular space (DTI-ALPS) is a diffusion-weighted MRI approach that proposes to evaluate the diffusion of CSF along perivascular spaces and is sensitive to aging effects and sleep physiology. The ALPS index is significantly lower in individuals with obstructive sleep apnea and N2 sleep stage duration in healthy older adults^37–39^. DTI-ALPS relies heavily on correctly identifying the orientation of perivascular spaces surrounding periventricular medullary veins, which is often difficult. Secondly, the sequence has a lower resolution on the order of 2mm^3^ isotropic and the signal in a normal DTI-sequence is not specific to CSF or ISF. Perivascular spaces are much smaller, and partial voluming is a major challenge in this approach. It is difficult to determine whether changes in the ALPS index are due to changes in fluid and/or fluid motion in perivascular compartments, to differences in blood signal, or frank axonal injury or degeneration in the vicinity of these vessels. IVIM also suffers from poorer resolution, but the multiple b-values allow discrimination between water pools with different mobility. Moreover, our ROIs in SAS can be placed more efficiently and reliably than identifying PVSs in a diffusion-weighted imaging sequence.

Another approach is to capture fluctuations in the BOLD fMRI signal within the ventricular CSF compartment or the use of ultrafast fMRI. Fultz et al. reported that BOLD fMRI signal in the 4^th^ ventricle was closely coupled with sleep stages. They and Helakari et al.^40^ further show that CSF fluctuations are associated with changes in CBF and cerebral blood volume (CBV), due to cardiac and respiratory fluctuations during sleep. These findings suggest that CSF dynamics within the ventricles are driven at least in part by cerebral hemodynamics. While the advantage of ASL is that it provides quantitative perfusion maps and, in our study, also a measure of trans-vascular permeability, the BOLD fMRI signal is faster, allowing better tracking of hemodynamics dependent on neuronal activity. Therefore, a fast fMRI scan would provide complementary information to ASL and should be considered in future multi-modal MRI studies.

There are several additional considerations for future studies. First, our measurements were made during awake states; thus, our findings represent the *result* of sleep and sleep deprivation. To define the dynamics of fluid transport through the course of physiological sleep, these methods should be evaluated during sleep and wake with in-magnet sleep-EEG studies. This would also improve our understanding of how the IVIM and ASL parameters are related to sleep. Finally, cognitive tests in this study were added to determine normal cognition but were not sensitive to sleep-related attention deficits. The inclusion of sleep-sensitive cognitive measures, such as a psychomotor vigilance task, could therefore aid in defining the relationship among sleep, cognitive performance, and MRI measures of glymphatic function in future studies.

## CONCLUSION

In this work, we showcase two brain imaging methods, IVIM and multi-echo, multi-delay ASL, to test their sensitivity to fluid transport changes upon sleep modulation and aging, seeking to define their potential utility in understanding glymphatic transport in the human brain. We observed that the diffusion-based IVIM approach was more sensitive to sleep physiology but not aging, while perfusion-based ASL approach was more sensitive to aging but not sleep physiology. Due to the clinical associations among aging, sleep disruption, glymphatic transport, and risk for dementia, both approaches are attractive to better understand the interaction between the vascular and CSF circulation and ISF solute transport during sleep and to assess the risk of dementia. Future studies could include measures of sleep architecture, markers of dementia pathogenesis, and sleep-sensitive cognitive measures in addition to these MRI measures.

## Abbreviations

ADRC: Alzheimer’s disease research center
AQP4: Aquaporin 4
ASL: arterial spin labeling
BOLD: blood oxygen level dependent
CSF: cerebrospinal fluid
FLAIR: fluid attenuated inversion recovery
fMRI: functional magnetic resonance imaging
GBCA: Gadolinium-based contrast agent
GM: gray matter
GPCOG: general practitioner’s cognitive assessment
ISF: interstitial fluid
IVIM: intravoxel incoherent motion
MIS: memory impairment screen
ROI: region of interest
SAS: subarachnoid space
SPLICE: Split acquisition of fast spin echo
WM: white matter

## ACKNOWLEDGEMENTS

This study was funded by the Royalty Research Fund at the University of Washington, awarded to Swati Rane Levendovszky. Swati Rane Levendovszky and Jeffrey Iliff were supported by the Medical Technology Enterprise Consortium (MTEC), Award: MT21006.129 and by R01 AG069960. Elaine Peskind is the Friends of Alzheimer’s Research Professor of Psychiatry and Behavioral Sciences at the University of Washington (UW) School of Medicine

## Author contributions

SRL: Study design, implementation, data analysis, manuscript writing. JF: Participant recruitment and data collection, EP: study design, data interpretation, manuscript editing, LV: implementation of multi-echo, multi-delay ASL protocol, data interpretation, manuscript editing, MJPvO: implementation of multi-echo, multi-delay ASL protocol, data interpretation, manuscript editing, JI: study design, data interpretation, manuscript writing and editing.

## REFERENCES

1. Iliff JJ, Lee H, Yu M, et al. Brain-wide pathway for waste clearance captured by contrast-enhanced MRI. J Clin Invest. 2013;123(3):1299–1309. doi:10.1172/JCI67677

2. Iliff JJ, Wang M, Liao Y, et al. A Paravascular Pathway Facilitates CSF Flow Through the Brain Parenchyma and the Clearance of Interstitial Solutes, Including Amyloid β. Sci Transl Med. 2012;4(147):147ra111. doi:10.1126/scitranslmed.3003748

3. Xie L, Kang H, Xu Q, et al. Sleep Drives Metabolite Clearance from the Adult Brain. Science. 2013;342(6156):373-377. doi:10.1126/science.1241224

4. Simon M, Wang MX, Ismail O, et al. Loss of perivascular aquaporin-4 localization impairs glymphatic exchange and promotes amyloid β plaque formation in mice. Alzheimers Res Ther. 2022;14(1):59. doi:10.1186/s13195-022-00999-5

5. Iliff JJ, Chen MJ, Plog BA, et al. Impairment of glymphatic pathway function promotes tau pathology after traumatic brain injury. J Neurosci. 2014;34(49):16180–16193. doi:10.1523/JNEUROSCI.3020-14.2014

6. Ishida K, Yamada K, Nishiyama R, et al. Glymphatic system clears extracellular tau and protects from tau aggregation and neurodegeneration. J Exp Med. 2022;219(3):e20211275. doi:10.1084/jem.20211275

7. Cui H, Wang W, Zheng X, et al. Decreased AQP4 Expression Aggravates L-Synuclein Pathology in Parkinson’s Disease Mice, Possibly via Impaired Glymphatic Clearance. J Mol Neurosci. 2021;71(12):2500–2513. doi:10.1007/s12031-021-01836-4

8. Zhang Y, Zhang C, He XZ, et al. Interaction Between the Glymphatic System and α- Synuclein in Parkinson’s Disease. Mol Neurobiol. 2023;60(4):2209–2222. doi:10.1007/s12035-023-03212-2

9. Ringstad G, Vatnehol SAS, Eide PK. Glymphatic MRI in idiopathic normal pressure hydrocephalus. Brain. 2017;140(10):2691–2705. doi:10.1093/brain/awx191

10. Watts R, Steinklein JM, Waldman L, Zhou X, Filippi CG. Measuring Glymphatic Flow in Man Using Quantitative Contrast-Enhanced MRI. AJNR Am J Neuroradiol. 2019;40(4):648–651. doi:10.3174/ajnr.A5931

11. Eide PK, Vinje V, Pripp AH, Mardal KA, Ringstad G. Sleep deprivation impairs molecular clearance from the human brain. Brain. 2021;144(3):863–874. doi:10.1093/brain/awaa443

12. Richmond SB, Rane S, Hanson MR, et al. Quantification approaches for magnetic resonance imaging following intravenous gadolinium injection: A window into brain-wide glymphatic function. Eur J Neurosci. Published online March 25, 2023. doi:10.1111/ejn.15974

13. Fultz NE, Bonmassar G, Setsompop K, et al. Coupled electrophysiological, hemodynamic, and cerebrospinal fluid oscillations in human sleep. Science. 2019;366(6465):628–631. doi:10.1126/science.aax5440

14. van Veluw SJ, Hou SS, Calvo-Rodriguez M, et al. Vasomotion as a Driving Force for Paravascular Clearance in the Awake Mouse Brain. Neuron. 2020;105(3):549–561.e5. doi:10.1016/j.neuron.2019.10.033

15. Williams SD, Setzer B, Fultz NE, et al. Neural activity induced by sensory stimulation can drive large-scale cerebrospinal fluid flow during wakefulness in humans. PLoS Biol. 2023;21(3):e3002035. doi:10.1371/journal.pbio.3002035

16. Le Bihan D, Breton E, Lallemand D, Aubin ML, Vignaud J, Laval-Jeantet M. Separation of diffusion and perfusion in intravoxel incoherent motion MR imaging. Radiology. 1988;168(2):497–505. doi:10.1148/radiology.168.2.3393671

17. Ohene Y, Harrison IF, Nahavandi P, et al. Non-invasive MRI of brain clearance pathways using multiple echo time arterial spin labelling: an aquaporin-4 study. Neuroimage. 2019;188:515–523. doi:10.1016/j.neuroimage.2018.12.026

18. Brodaty H, Pond D, Kemp NM, et al. The GPCOG: a new screening test for dementia designed for general practice. J Am Geriatr Soc. 2002;50(3):530–534. doi:10.1046/j.1532-5415.2002.50122.x

19. Buschke H, Kuslansky G, Katz M, et al. Screening for dementia with the memory impairment screen. Neurology. 1999;52(2):231–238. doi:10.1212/wnl.52.2.231

20. Schick F. SPLICE: sub-second diffusion-sensitive MR imaging using a modified fast spin-echo acquisition mode. Magn Reson Med. 1997;38(4):638–644. doi:10.1002/mrm.1910380418

21. Petitclerc L, Schmid S, Hirschler L, van Osch MJP. Combining T2 measurements and crusher gradients into a single ASL sequence for comparison of the measurement of water transport across the blood–brain barrier. Magnetic Resonance in Medicine. 2021;85(5):2649–2660. doi:10.1002/mrm.28613

22. Teeuwisse WM, Schmid S, Ghariq E, Veer IM, van Osch MJP. Time-encoded pseudocontinuous arterial spin labeling: basic properties and timing strategies for human applications. Magn Reson Med. 2014;72(6):1712–1722. doi:10.1002/mrm.25083

23. Jenkinson M, Beckmann CF, Behrens TEJ, Woolrich MW, Smith SM. FSL. NeuroImage. 2012;62(2):782-790. doi:10.1016/j.neuroimage.2011.09.015

24. Killer HE, Laeng HR, Flammer J, Groscurth P. Architecture of arachnoid trabeculae, pillars, and septa in the subarachnoid space of the human optic nerve: anatomy and clinical considerations. Br J Ophthalmol. 2003;87(6):777–781. doi:10.1136/bjo.87.6.777

25. Saboori P, Sadegh A. Histology and Morphology of the Brain Subarachnoid Trabeculae. Anat Res Int. 2015;2015:279814. doi:10.1155/2015/279814

26. Harrison IF, Siow B, Akilo AB, et al. Non-invasive imaging of CSF-mediated brain clearance pathways via assessment of perivascular fluid movement with diffusion tensor MRI. Elife. 2018;7:e34028. doi:10.7554/eLife.34028

27. Le Bihan D. What can we see with IVIM MRI? Neuroimage. 2019;187:56–67. doi:10.1016/j.neuroimage.2017.12.062

28. Pérès EA, Etienne O, Grigis A, Boumezbeur F, Boussin FD, Le Bihan D. Longitudinal Study of Irradiation-Induced Brain Microstructural Alterations With S-Index, a Diffusion MRI Biomarker, and MR Spectroscopy. Int J Radiat Oncol Biol Phys. 2018;102(4):1244–1254. doi:10.1016/j.ijrobp.2018.01.070

29. Chen JJ, Rosas HD, Salat DH. Age-Associated Reductions in Cerebral Blood Flow Are Independent from Regional Atrophy. Neuroimage. 2011;55(2):468–478. doi:10.1016/j.neuroimage.2010.12.032

30. Tarumi T, Zhang R. Cerebral Blood Flow in Normal Aging Adults: Cardiovascular Determinants, Clinical Implications, and Aerobic Fitness. J Neurochem. 2018;144(5):595–608. doi:10.1111/jnc.14234

31. Alosco ML, Brickman AM, Spitznagel MB, et al. Reduced cerebral blood flow and white matter hyperintensities predict poor sleep in heart failure. Behav Brain Funct. 2013;9:42. doi:10.1186/1744-9081-9-42

32. Tüshaus L, Omlin X, Tuura RO, et al. In human non-REM sleep, more slow-wave activity leads to less blood flow in the prefrontal cortex. Sci Rep. 2017;7(1):14993. doi:10.1038/s41598-017-12890-7

33. Mahroo A, Konstandin S, Günther M. Blood–Brain Barrier Permeability to Water Measured Using Multiple Echo Time Arterial Spin Labeling MRI in the Aging Human Brain. Journal of Magnetic Resonance Imaging. n/a(n/a). doi:10.1002/jmri.28874

34. Ford JN, Zhang Q, Sweeney EM, et al. Quantitative Water Permeability Mapping of Blood-Brain-Barrier Dysfunction in Aging. Front Aging Neurosci. 2022;14:867452. doi:10.3389/fnagi.2022.867452

35. Gold BT, Shao X, Sudduth TL, et al. Water exchange rate across the bloodLbrain barrier is associated with CSF amyloidLβ 42 in healthy older adults. Alzheimers Dement. 2021;17(12):2020–2029. doi:10.1002/alz.12357

36. Palomares JA, Tummala S, Wang DJJ, et al. Water Exchange across the Blood-Brain Barrier in Obstructive Sleep Apnea: An MRI Diffusion-Weighted Pseudo-Continuous Arterial Spin Labeling Study. J Neuroimaging. 2015;25(6):900–905. doi:10.1111/jon.12288

37. Hsiao WC, Chang HI, Hsu SW, et al. Association of Cognition and Brain Reserve in Aging and Glymphatic Function Using Diffusion Tensor Image-along the Perivascular Space (DTI-ALPS). Neuroscience. Published online April 6, 2023:S0306–4522(23)00163-X. doi:10.1016/j.neuroscience.2023.04.004

38. Lee HJ, Lee DA, Shin KJ, Park KM. Glymphatic system dysfunction in obstructive sleep apnea evidenced by DTI-ALPS. Sleep Med. 2022;89:176–181. doi:10.1016/j.sleep.2021.12.013

39. Taoka T, Masutani Y, Kawai H, et al. Evaluation of glymphatic system activity with the diffusion MR technique: diffusion tensor image analysis along the perivascular space (DTI-ALPS) in Alzheimer’s disease cases. Jpn J Radiol. 2017;35(4):172–178. doi:10.1007/s11604-017-0617-z

40. Helakari H, Korhonen V, Holst SC, et al. Human NREM Sleep Promotes Brain-Wide Vasomotor and Respiratory Pulsations. J Neurosci. 2022;42(12):2503–2515. doi:10.1523/JNEUROSCI.0934-21.2022

