## Supplementary figure for "Preliminary cross-sectional investigations into the human glymphatic system using multiple novel non-contrast MRI methods"

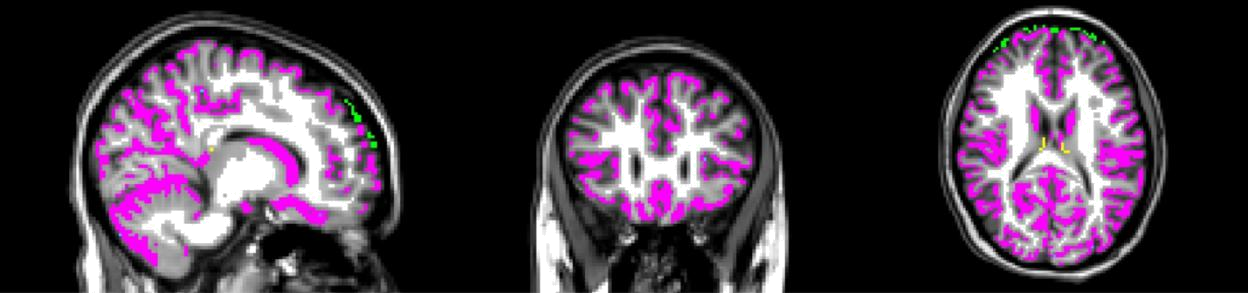

**Supplementary Figure 1:** ROI generation for the subarachnoid space (SAS, green) and choroid

plexus (CP, yellow), gray matter (GM, magenta), and white matter (WM, white).
